# The Interplay Between Cell-cell Signaling and Negative Feedback Reduces Population Noise While Enhancing Single-cell Variability

**DOI:** 10.1101/2025.01.07.631833

**Authors:** Xinying Ren

## Abstract

Negative feedback is a well-known mechanism for attenuating noise and enhancing robustness in biological systems. When coupled with cell-cell signaling, it provides a strategy for achieving population-level control in multicellular systems. While cell-cell signaling alone tends to reduce cell-to-cell variability by averaging fluctuations across cells, its interplay with negative feedback can produce contrasting effects on noise regulation at single-cell and population levels. Therefore, to design population-level controllers that achieve robust behaviors with attenuated noise, a systematically understanding of how different noise sources impact cell gene expressions at these levels become a critical challenge. Here, we investigate noise regulation in a quorum sensing-based negative feedback system, focusing on two extrinsic noise sources: process noise from target gene dynamics and measurement noise from quorum sensing dynamics. Our results reveal that signal-based negative feedback significantly reduces process noise at the population level, especially for dynamic noise. However, at the single-cell level, it enhances variability, with increased noise levels under faster signal diffusion and higher population density. In contrast, measurement noise is consistently attenuated at both single-cell and population levels through the coupled cell-cell signaling and negative feedback, under conditions of faster diffusion and higher density.

## I. Introduction

Noise is ubiquitous in cellular processes, potentially disrupting stable and robust cellular functions and compromising homeostasis in multicellular systems [1]. Cells have evolved various noise-attenuation strategies at the single-cell level, for example, reducing translation burst rates [2], incorporating intracellular negative feedback regulation [3], [4], and employing ultrasensitive switches and feedforward loops [5]–[7].

In multicellular systems, individual cells aggregate to form populations and communities. While single-cell level noise-attenuation strategies are essential, cell-cell signaling has emerged as a major mechanism that promotes population-level robustness [8], [9]. Signaling mechanisms facilitate synchronization and consensus across the population, protecting cells against noise via averaging effects and cooperative interactions [10]. In quorum sensing (QS) systems, for instance, extrinsic noise is reduced through “diffusional dissipation”, where rapid diffusion of signaling molecules and fast turnover of transcriptional regulators dampen noise in target gene expression [11].

By integrating cell-cell signaling with intracellular negative feedback, population-level feedback has shown significant potential to achieve robust gene expression in multicellular systems [12]–[14]. Negative feedback mechanisms within each cell work alongside these signaling pathways to dampen fluctuations, allowing the overall population to achieve a more stable expression profile. However, this approach does not explicitly identify which sources of noise or disturbances are attenuated within the closed-loop system, as many studies examine combined intrinsic and extrinsic noise from multiple intracellular and extracellular sources. Additionally, it remains unclear if single-cell variability and population-level expression fluctuations are reduced in the same manner and simultaneously by cell-cell signaling. As a result, current designs for population-level control lack a systematic framework to optimize noise reduction and disturbance rejection performances for distinct noise sources. Furthermore, design principles for the independent control of single-cell variability and population robustness are still unclear.

In this paper, we address these gaps by analyzing noise regulation in a quorum sensing-based population feedback system (Section II). We focus on two extrinsic noise sources: process noise, originating from heterogeneous expression of the target gene expression, and measurement noise, arising from fluctuations in quorum sensing dynamics, as shown in Fig.1A. Using an ordinary differential equation (ODE) model, we derive the mean and variance of gene expression at steady-state at both single-cell and population levels. Through frequency analysis and power spectrum density (PSD) calculations, we investigate how process and measurement noise propagate differently in the closed-loop system (Section III, IV). Our findings are further validated by simulations of single-cell trajectories in populations under open-loop, intracellular feedback and population-level feedback conditions. Finally, we discuss design principles for population-level feedback to control gene expression variance and population robustness (Section V).

**Fig. 1.**
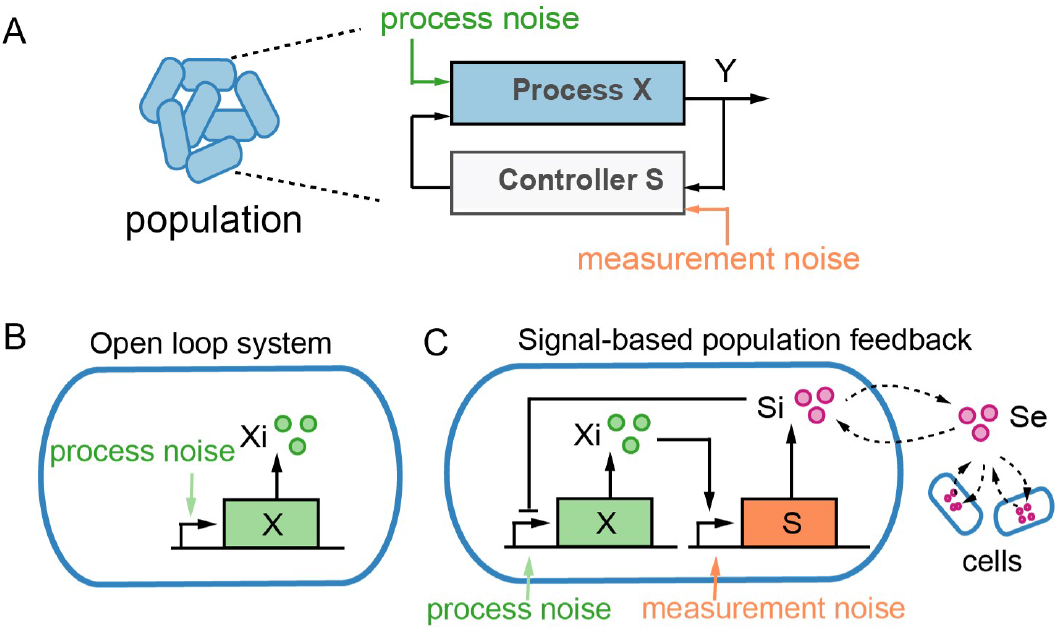
Noise regulation in population-level feedback systems. (A) Block diagram of a closed-loop feedback system. Process noise arises in the process dynamics, and measurement noise is introduced in the controller dynamics. (B) Schematic of an open-loop system of the target gene expression. Process noise originates from the varying transcriptional dynamics of the target gene *X*. (C) Schematic of a signal-based population feedback system. A quorum sensing signaling molecule, *S*, diffuses freely between intracellular and extracellular environments. The signaling molecule collectively senses and represses *X* production across individual cells.

## II. System Descriptions and Mathematical Models

### A. Open-loop system

Our control objective is to regulate the expression of a target gene *X* in a cell population. Without feedback, *X* undergoes constant production and cell dilution, as illustrated in Fig.1B. Let *x*_*i*_ denote the expression level (protein concentration) of *X* in the *i*th cell, and *y* denote the population-level expression *Y*, defined as the sum of *X* expression across all cells. For a population of *n* cells, the cellular dynamics can be modeled by the following ODE, for *i* = 1, 2, …, *n*:

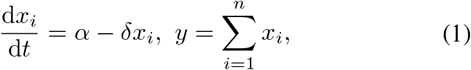

where *α* and *δ* are the production and dilution rates, respectively. The steady-state solutions are solved as:

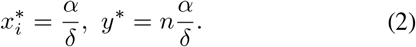

### B. Signal-based population feedback system

To incorporate population-level feedback, we introduce a quorum sensing system where a signaling molecule, *S*, acts as a negative feedback controller. As shown in Fig.1C, *X* is a transcriptional factor that activates the synthesis of *S*, and in turn, *S* represses the transcription of *X*. The signaling molecule *S* diffuses freely across cell membranes, with a constant dilution in the extracellular environment.

Let *s*_*i*_ denote the intracellular concentration of *S* in the *i*th cell, and *s*_*e*_ denote the extracellular concentration. The dynamics of the closed-loop system are modeled as:

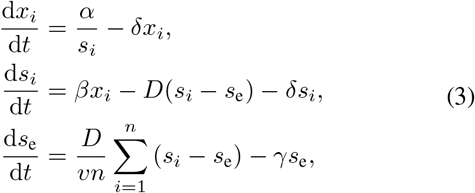

where *β* and *γ* are the production rate and extracellular dilution rate of *S*, respectively. The parameter *D* is the diffusion rate of *S* across cell membranes, and *v* is the ratio of extracellular volume *V*_*e*_ to total cell volume, defined as 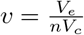, where *V*_*c*_ is the volume of a single cell. The steady-state solutions for this system are:

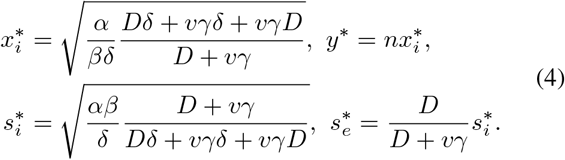

Here we analyze two specific conditions:

#### 1) No diffusion (D = 0)

In this case, the signaling molecule *S* remains intracellular, providing negative feedback only within individual cells. Since there is no cell-cell signaling, it is equivalent to intracellular-only negative feedback. The system’s steady-state solutions become:

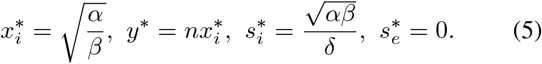

#### 2) Fast diffusion (D → ∞)

When diffusion is rapid, intracellular and extracellular concentrations of *S* reach an equilibrium almost instantaneously. The steady-state solutions become:

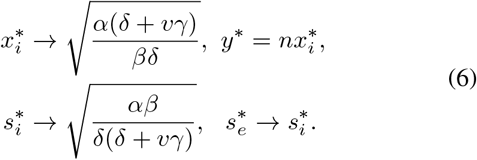

In quorum sensing systems of free and diffusible signaling molecules, the diffusion rate *D* depends on the viscosity of the environment. In liquid environments, *D* is typically much larger than the rates of intracellular reactions, allowing *s*_*e*_ and *s*_*i*_ to reach a balanced concentration efficiently [15]. In other cell-cell signaling systems that involve transportation of signaling molecules, the diffusion rate *D* also depends on the transportation efficiency across cell membranes [16].

### C. Parameter values

To simplify analysis, we set *α* = 1, *δ* = 1 and *V*_*c*_ = 1. The parameter *β* is adjusted to maintain consistent steady-state levels across different systems. For the open-loop system, these parameter settings yield the following steady-state concentrations: 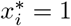 and *y*^*^ = *n*. For the feedback systems, *β* is set to:

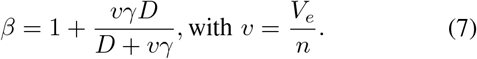

This parameter setting ensures the same steady-state concentrations of the target gene for all systems:

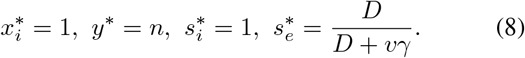

In the following sections, we explore how process and measurement noise propagates in different systems with the same steady state *x*_*i*_ = 1, *y* = *n*. We consider three conditions: the open-loop system, the intracellular-only negative feedback system (*D* = 0), and the population-level feedback system (*D >* 0). Specifically, we examine the effects of several parameters on noise reduction and disturbance rejection performances at single-cell and population levels, including diffusion rate *D*, extracellular dilution rate *γ*, population density *n*, and extracellular volume *V*_*e*_.

## III. Process Noise Reduction at Single-cell and Population Levels

We first study process noise, which arises from transcriptional variations of *X* within individual cells. The noise in the *X* production rate in the *i*th cell is denoted by Δ*α*_*i*_. Using the Langevin approach with linearized ODEs around the steady state, we calculate the noise statistics for *x*_*i*_ (single-cell level) and *y* (population level). First, we model the process noise Δ*α*_*i*_ as static, independent and identically distributed (i.i.d.) random variables. Then we extend this model to dynamic process noise, Δ*α*_*i*_(*t*), represented as stochastic processes, and evaluate the power spectrum density (PSD) in the frequency domain.

The linearized open-loop system is given by the following equation:

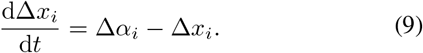

For the intracellular negative feedback system, the linearized ODEs are:

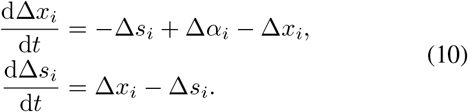

For the signal-based population feedback system, we have:

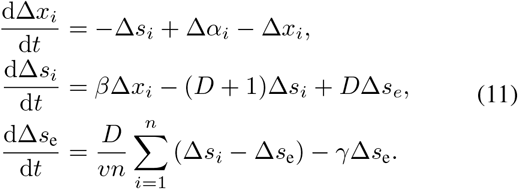

### A. *Static noise* Δ*α*_*i*_

Assuming static, i.i.d. process noise Δ*α*_*i*_∼ 𝒩 (0, *σ*^2^), we solve for the steady-state solutions for these systems.

#### 1) Open-loop system

From Eq (9), we obtain:

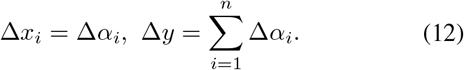

The noise statistics are:

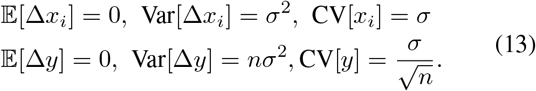

Without feedback, process noise persists in expression levels at both single-cell and population levels.

#### 2) Intracellular negative feedback system

From Eq (10), we obtain:

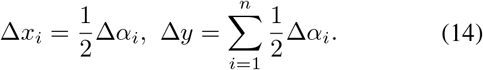

The noise statistics are:

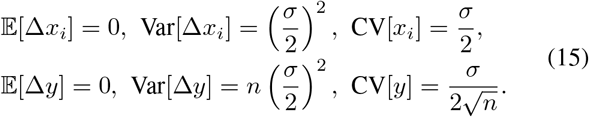

Intracellular negative feedback attenuates process noise within each cell independently. At both single-cell and population levels, it reduces the coefficient of variation (CV) by half.

#### 3) Signal-based population negative feedback system

Solving Eq (11) yields the following expression for single-cell level noise, which depends on process noise from each individual cell and other neighboring cells:

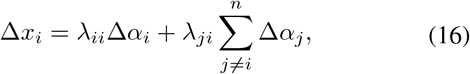

where

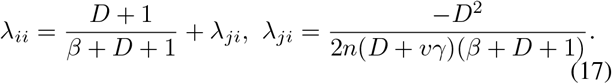

The characteristics of noise in *X*_*i*_ are:

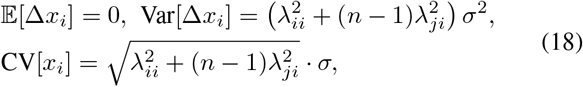

depending on parameters *D, γ, n* and *V*_*e*_.

To gain an insight on how the interplay between cell-cell signaling and negative feedback affects expression noise at single-cell level, we consider a fast diffusion scenario. We assume *D* ≫ 1, *vγ*, approximating *β* ≈1 + *vγ*. Then we can approximate:

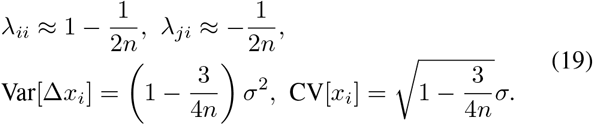

In this case, 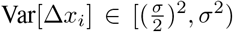. For a small population with only one single cell (*n* = 1), noise is minimized with variance value 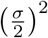. As *n* increases, Var[Δ*x*_*i*_] increases, approaching its maximum *σ*^2^. Compared with intracellular-only negative feedback, population-level feedback is less effective at single-cell level noise reduction when diffusion is fast.

To further explore how Var[Δ*x*_*i*_] depends on the parameters *D, γ, n* and *V*_*e*_, we numerically solve for Var[Δ*x*_*i*_] and present the results in Fig.2A and B. At fixed population density *n* and extracellular volume *V*_*e*_, process noise in *X*_*i*_ increases with diffusion rate *D* but decreases with extracellular dilution rate *γ*. Meanwhile, for fixed *D* and *γ*, process noise in *X*_*i*_ increases with *n* and decreases with *V*_*e*_. These results indicate that single-cell variability is enhanced under faster signaling diffusion in populations with higher density.

**Fig. 2.**
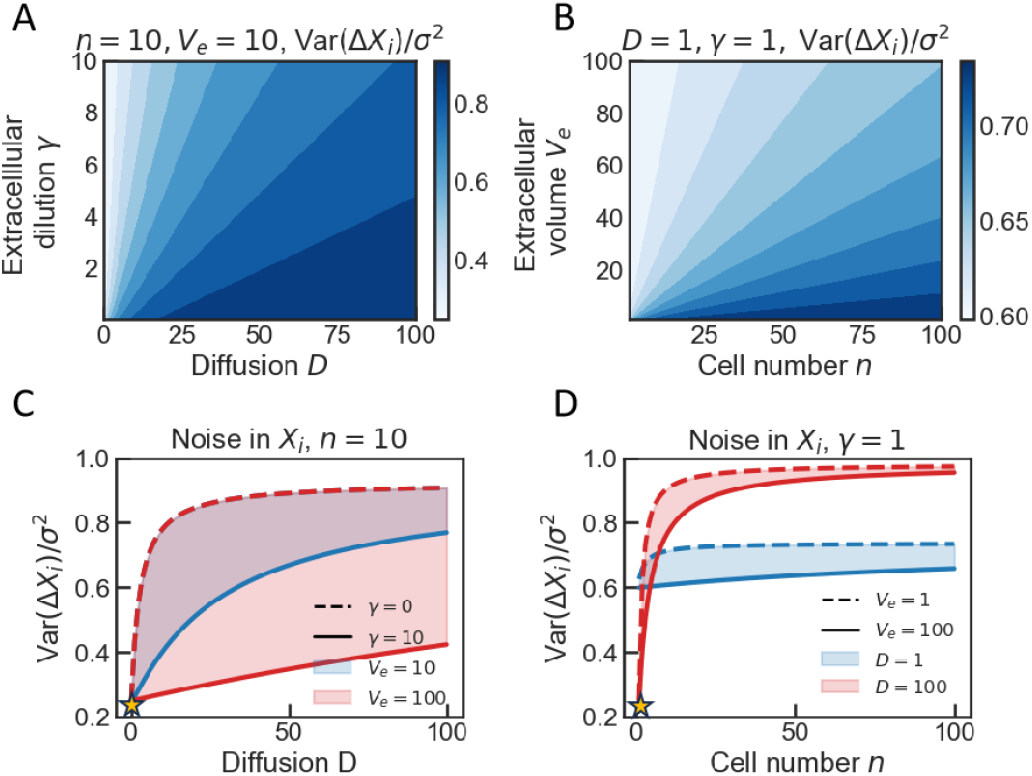
Numerical solutions for the static noise variance at the single-cell level under signal-based population feedback. (A) Dependence of Var[Δ*x*_*i*_] on diffusion rate *D* and extracellular dilution rate *γ*, given small population density and extracellular volume. (B) Dependence of Var[Δ*x*_*i*_] on population density *n* and extracellular volume *V*_*e*_, with small signal diffusion and dilution rates. (C) Dependence of Var[Δ*x*_*i*_] on the dominant parameter *D*. The minimum noise variance 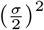 is achieved when *D* = 0, marked by the yellow star. Shaded regions between dashed and solid lines represent variance regimes across a range of *γ* values, with red and blue areas corresponding to two different conditions of *V*_*e*_. (D) Dependence of Var[Δ*x*_*i*_] on the dominant parameter *n*. The minimum noise variance 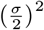 is achieved when *n* = 1 and *D* is large, marked by the yellow star. Shaded regions between dashed and solid lines represent variance regimes across a range of *V*_*e*_ values, with red and blue areas corresponding to two different conditions of *D*.

Specifically, as shown in Fig.2C and D, when *D* → 0 or *D* → ∞, *n* → 1, the process noise at the single-cell level is significantly reduced, with Var[Δ*x*_*i*_] reaching its theoretical minimum:

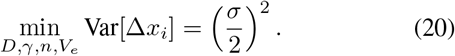

The first condition of no dilution (*D* →− 0), aligns with the analytical result for intracellular-only feedback system, as shown in Eq (15). The second condition (*D* → ∞, *n* → 1) represents fast diffusion but with a small cell population, which is consistent with the analytical result in Eq (19). Both conditions indicate that the noise is reduced to its minimum when there is minimal averaging effects from cell-cell signaling either due to no diffusion or low population density. In contrast to the understanding that cell-cell signaling, by itself, tends to reduce cellular variability by averaging fluctuations across cells, the interplay with negative feedback induces an opposite effect.

This analysis implies that cell-cell signaling can diminish the effectiveness of process noise reduction in systems governed by population-level negative feedback. In fact, when the averaging effect from cell-cell signaling is strong (*D* and *n* are large), the intracellular concentration of signaling molecules becomes effectively noise-free, i.e., Var[Δ*s*_*i*_] → 0. As a result, *S* no longer induces the compensatory regulation in *X*_*i*_, as it would in intracellular-only feedback. The process noise is therefore sustained, leading to high single-cell variability.

Detailed analysis and computation on the variance of *S* are provided in Appendix.

However, at the population level, we find:

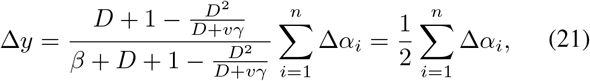

yielding:

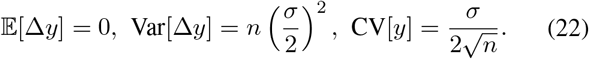

The population-level expression *Y* exhibits the same noise variance as observed in the intracellular-only feedback system, shown in Eq (15). It indicates that while single-cell level noise remains high, population-level fluctuations are substantially reduced. This attenuation of process noise is robust to variations in other parameters *D, γ* and *V*_*e*_.

To further investigate why population-level feedback leads to large variance at the single-cell level but small variance at the population level, we directly derive Var[Δ*y*] from Var[Δ*x*_*i*_] as follows:

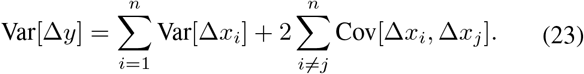

Based on Eq (16) and (17), we solve for the covariance:

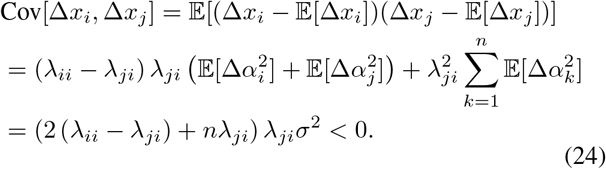

The negative covariance indicates that cell-cell signaling introduces a negative correlation between noise in individual cells. This mechanism allows high single-cell variability, while the collective noise at the population level is effectively minimized.

### B. Simulation of single-cell and population trajectories

To further validate our theoretical analysis, we run simulations of single-cell and population-level expression trajectories in the presence of static noise Δ*α*_*i*_, and compare steady-state noise distributions across open-loop and feedback systems. We set *σ* = 0.2, thus in the simulation, *X* production rate in the *i* the cell is *α*_*i*_ = 1 + Δ*α*_*i*_, following 𝒩 (1, 0.2^2^). We assume that the extracellular volume equals to the total cell volume, thus we set *v* = 1.

First, we examine single-cell level noise in a population of *n* = 1000 cells. As shown in Fig.3A and B, when *D* is large, we find that *X*_*i*_ concentration trajectories exhibit a larger variance compared to *S*_*i*_ concentration. This simulation result confirms that strong averaging effects from cell-cell signaling weakens noise attenuation efficiency of the negative feedback. From the steady-state distribution, we compute CV[*x*_*i*_] ≈ *σ*, which agrees with the analytical solution in Eq (19).

**Fig. 3.**
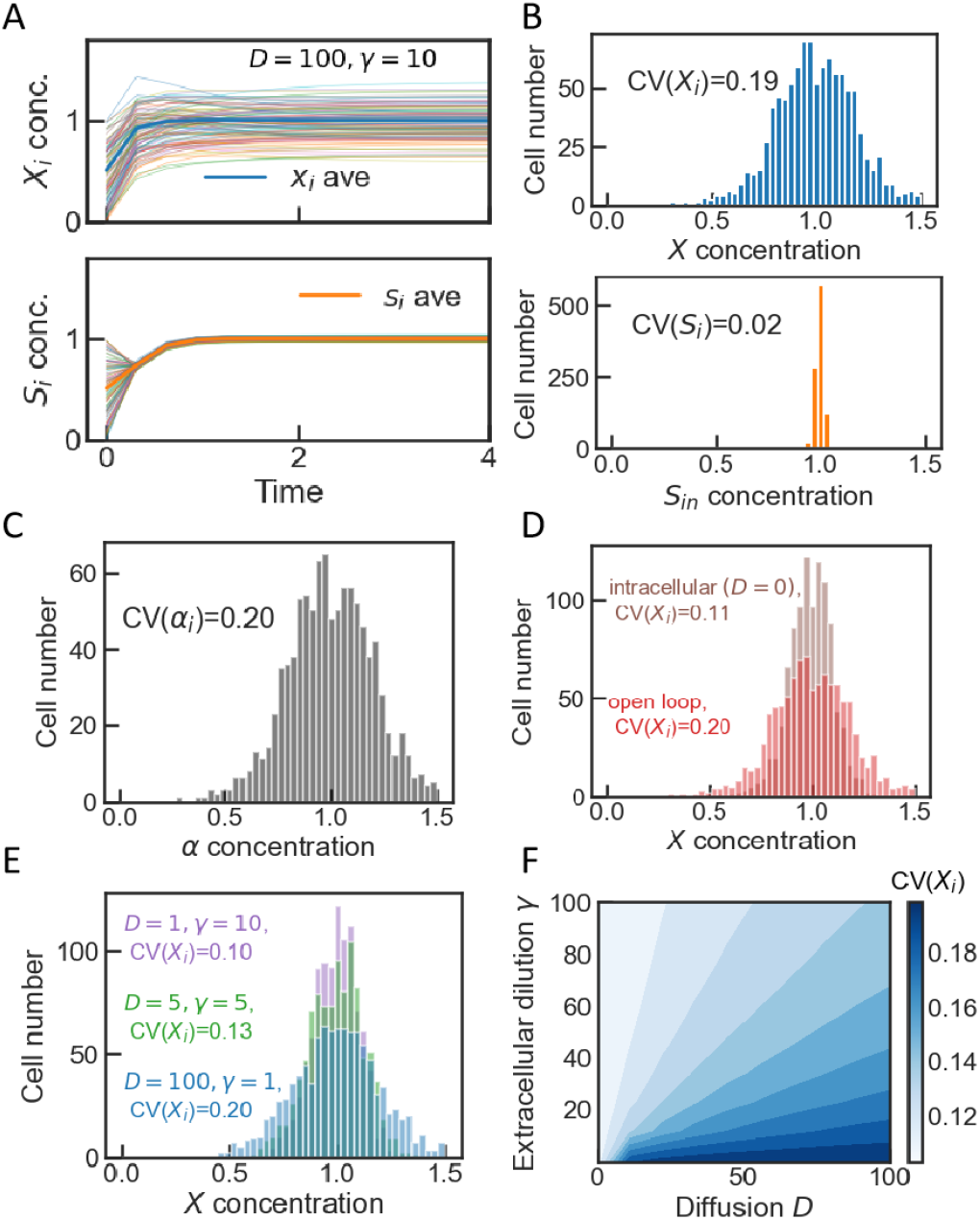
Simulation results of single-cell expression trajectories and steady-state distributions with static process noise Δ*α*_*i*_. (A) Sampled trajectories (100 out of *n* =1000 cells) of target gene expression and intracellular signaling molecule concentrations under a fast diffusion and extracellular dilution condition (*D* = 100, *γ* = 10, *n* = 1000, *V*_*e*_ = 1000). (B) Steady-state distributions and corresponding CV values of *X*_*i*_ and *S*_*i*_, where 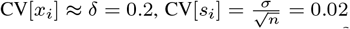. (C) Distribution of *X* production rate *α*_*i*_ with process noise Δ*α*_*i*_ ∼ 𝒩 (0, *σ*^2^), where *σ* = 0.2. (D) Steady state distributions of *X*_*i*_ in the open-loop system (red) and the intracellular negative feedback system (brown). (E) Steady state distribution of *X*_*i*_ in the signal-based population feedback system with different diffusion and extracellular dilution rates. (F) Dependence of CV[*x*_*i*_] on *D* and *γ*.

To compare noise reduction efficiency in different systems: open loop, intracellular negative feedback and signal-based population feedback, we present *α*_*i*_ distribution in Fig.3C and *X*_*i*_ distributions at steady state in Fig.3D and E. We find that the open-loop system maintains the full level of process noise, while intracellular negative feedback reduces the noise CV_in_[*x*_*i*_] by half. For signal-based population feedback, CV_s_[*x*_*i*_] is close to its minimum value 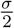 when *D* is small and *γ* is large. As *D* increases and *γ* decreases, CV_s_[*x*_*i*_] increases to its maximum *σ*, as shown in Fig.3F.

Next, we assess noise attenuation at the population level. We simulate 500 populations, each consisting of *n* = 100 cells. The trajectories of population-level expression of *Y* with population feedback are illustrated in Fig.4A. We present *α*_*i*_ distribution in Fig.4B and *Y* distributions of different systems at steady state in Fig.4C and D. Signal-based population feedback achieves a steady-state distribution with 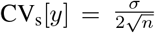, which is robust across varying values of *D* and *γ*. Similarly, intracellular negative feedback also reduces the noise in population-level expression with 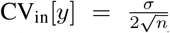. Meanwhile, the open-loop system maintains a large noise with 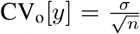. These simulation results agree with analytical solutions in Eq (13), (15) and (22), which all demonstrate that population feedback and intracellular feedback reduce static process noise at population-level equally effective.

**Fig. 4.**
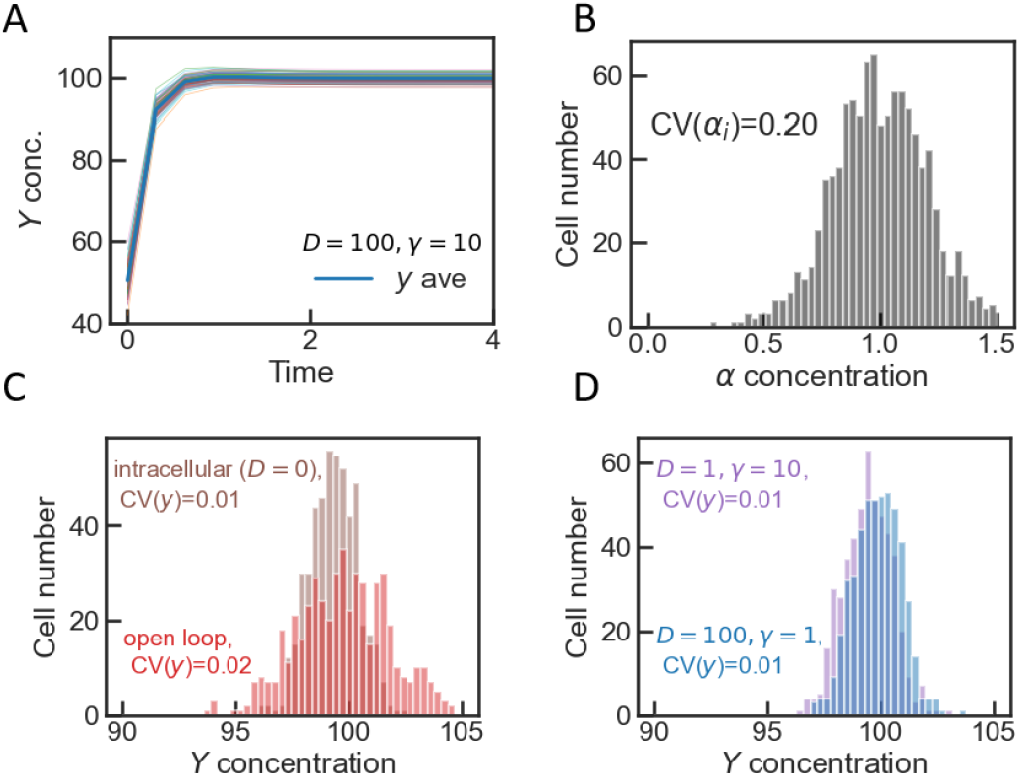
Simulation results of population-level expression trajectories and steady-state distributions with static process noise Δ*α*_*i*_. (A) Sampled trajectories (100 out of 500 populations) of total target expression under a fast diffusion condition (*D* = 100, *γ* = 1, *n* = 100, *V*_*e*_ = 100). (B) Distribution of *X* production rate *α*_*i*_ with noise Δ*α*_*i*_ ∼ 𝒩 (0, *σ*^2^), where *σ* = 0.2. (C) Steady state distributions of *Y* in the open-loop system (red) and the intracellular negative feedback system (brown). (D) Steady state distribution of *Y* in the signal-based population feedback system with different diffusion and extracellular dilution rates.

### C. Dynamic noise Δα_i_(t) and PSD calculation

While negative feedback attenuates noise at low frequencies,it can paradoxically amplify noise at higher frequencies [4]. In this section, we investigate how cell-cell signaling shapes the frequency-dependent noise reduction properties of negative feedback. We introduce dynamic process noise Δ*α*_*i*_(*t*), which is time-dependent and contains components across a range of frequencies, and calculate the power spectrum density (PSD).

We perform a Fourier transformation of the population feedback system as described in Eq (11). The resulting population-level noise is:

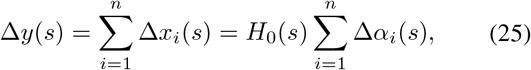

where the transfer function *H*_0_(*s*) is given by:

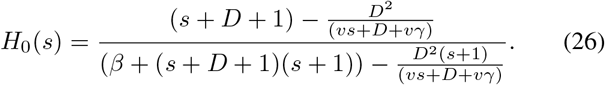

This describes how all process noise 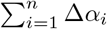 propagates to the population-level expression at frequency domain. At single-cell level, the expression noise is given by:

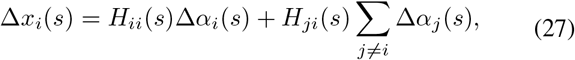

with the corresponding transfer functions *H*_*ii*_(*s*) and *H*_*ji*_(*s*) as follows:

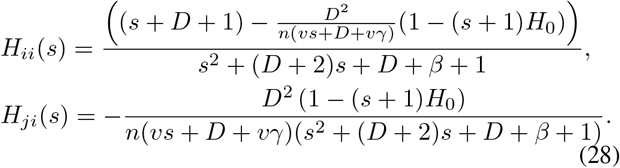

*H*_*ii*_(*s*) and *H*_*ji*_(*s*) describe how process noise from an individual cell and its neighboring cells affect *X* expression in the *i*th cell.

We present the Bode plot (magnitude) of the transfer function *H*_0_(*s*) in Fig.5A. Notably, compared with intracellular-only negative feedback, population-level feedback exhibits similar noise attenuation at low frequencies, which aligns with our analysis for the static noise scenario. However, in the high-frequency regime, cell-cell signaling facilitates further noise reduction rather than amplification. Specifically, when signal diffusion rate *D* and extracellular dilution rate *γ* are fast, noise attenuation in the high-frequency domain becomes more pronounced.

**Fig. 5.**
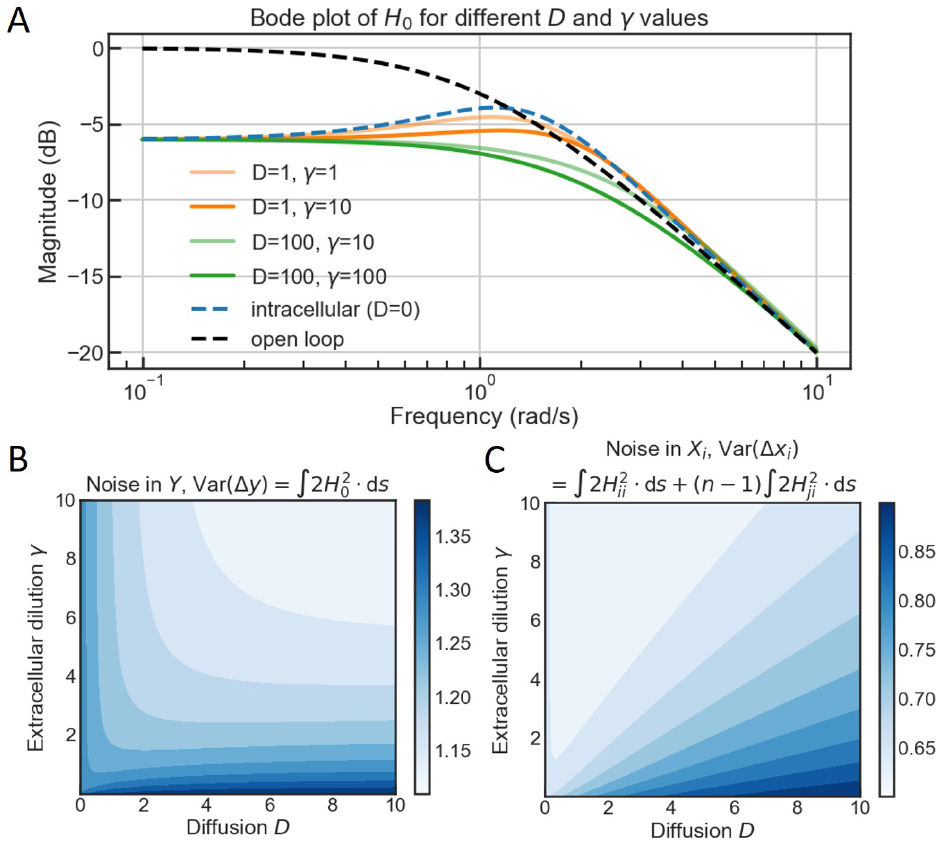
Numerical analysis of transfer functions in the frequency domain and expression variance under dynamic process noise Δ*α*_*i*_(*t*). (A) Bode plots (magnitude) of the transfer functions *H*_0_(*s*) for the open loop (dashed black), intracellular feedback (blue dashed) and signal-based population feedback (orange and green) with varying parameters. (B) Dependence of Var[Δ*y*] on diffusion *D* and extracellular dilution *γ*. (C) Dependence of Var[Δ*x*_*i*_] on diffusion *D* and extracellular dilution *γ*.

This observation suggests that the interplay between cell-cell signaling and negative feedback mostly enhances dynamic noise reduction at the population level. To evaluate noise reduction efficiency, we calculate the variance of Δ*y*. According to Plancherel’s theorem, variance in the temporal domain can be computed by integrating the PSD over the frequency domain. Assuming that the spectrum of Δ*α*_*i*_ is white with an effective magnitude 1, the PSD of noise in *Y* and the variance are calculated as follows:

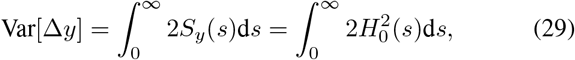

given that the PSD of noise in *Y*, denoted by *S*_*y*_(*s*), depends on the PSD of noise in all *α*_*i*_, denoted by 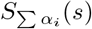:

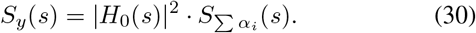

We compute Var[Δ*y*] for different parameter values of *D* and *γ*, and present the results in Fig.5B. It shows that, at the population level, dynamic process noise reduction efficiency improves with increasing diffusion and extracellular dilution rates.

Furthermore, we examine the single-cell level expression noise by calculating the variance from the PSD of noise in *X*_*i*_ in a similar way, and plot Var[Δ*x*_*i*_] in Fig.5C. The dependence of *D* and *γ* is consistent with our analysis of static noise regulation at single-cell level. However, it shows an opposite trend compared to the result at population level. Specifically, increasing signal diffusion rate *D* leads to higher cell-to-cell variability while it reduces population-level fluctuations.

## IV. Measurement Noise Reduction at Single-cell and Population Levels

In this section, we examine measurement noise, which arises from cell-cell signaling dynamics. Cell-cell signaling mechanisms, for example, quorum sensing, involve complicated dynamics including the synthesis, receptor binding and degradation of signaling molecules. Thus, fluctuations in the cell-cell signaling pathway can introduce noise in target gene expression.

Here, we consider a measurement noise as fluctuations in the production rate of signaling molecules in all cells, denoted by Δ*β*_*i*_. Following a similar approach as for process noise, we analytically derive for the mean, variance and coefficient of variation (CV) at steady state for static noise, verify results through simulations, and compute noise statistics in dynamic settings. Detailed modeling, derivation and results are provided in Appendix.

### A. *Static noise* Δ*β*_*i*_

For intracellular negative feedback (*D* = 0), we obtain:

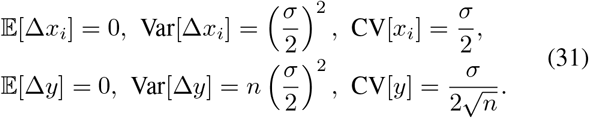

This result shows that measurement noise is attenuated in the same manner as process noise at both single-cell and population levels, as described in Eq (15).

For signal-based population negative feedback, we have:

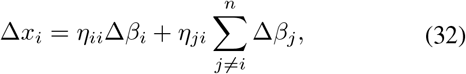

where

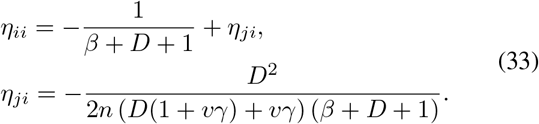

If we consider a fast diffusion condition where *D* ≫ 1, *vγ*, we can approximate the statistics of noise in *X*_*i*_ as:

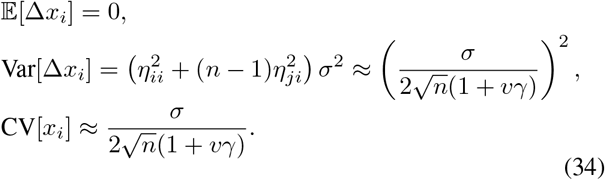

In this scenario, Var[Δ*x*_*i*_] decreases with population density *n*, extracellular volume *V*_*e*_ and dilution *γ*, approaching the minimum value of 0. This indicates that measurement noise is effectively attenuated by strong averaging effects from cell-cell signaling, which reduces the variability in single cells. The noise is completely diminished when cell-cell signaling is extremely strong (*D, n* → ∞).

At the population level, we derive:

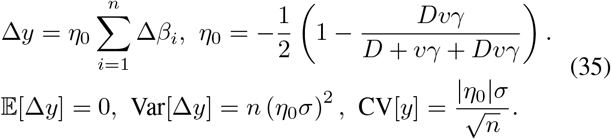

Note that *η*_0_ *<* 0. When *D, vγ* ≫ 1, *η*_0_ approaches its maximum value 0. When *D* →− 0 or *vγ* →− 0, *η*_0_ approaches its minimum value 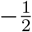. Given that 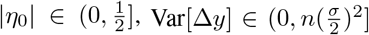. This indicates that signal-based population feedback reduces measurement noise at the population level much more effectively than intracellular-only negative feedback. As diffusion and extracellular dilution rates increase, Var[Δ*y*] decreases, leading to a population-level expression with Var[Δ*y*] → 0.

The analysis for static measurement noise shows that cell-cell signaling significantly improves noise reduction and disturbance rejection performances at both single-cell and population levels, which is consistent with previous studies suggesting that strong averaging effects among cells help attenuating fluctuations [11]–[13]. The reduction efficiency is enhanced with higher diffusion rate *D* and extracellular dilution rate *γ*, and larger population density *n* and extracellular volume *V*_*e*_.

### B. Dynamic noise Δβ_i_(t) and PSD calculation

We further analyze measurement noise in the frequency domain and calculate the variance of noise in *X*_*i*_ and *Y* from their PSD. The result is consistent with our analysis for static measurement noise. As shown in the Bode plot in Fig.6A, signal-based population feedback attenuates measurement noise at all frequencies. Both Var[Δ*x*_*i*_] and Var[Δ*y*] decrease with *D* and *γ*, as shown in Fig.6B and C, suggesting that strong cell-cell signaling and averaging effects improve noise attenuation.

**Fig. 6.**
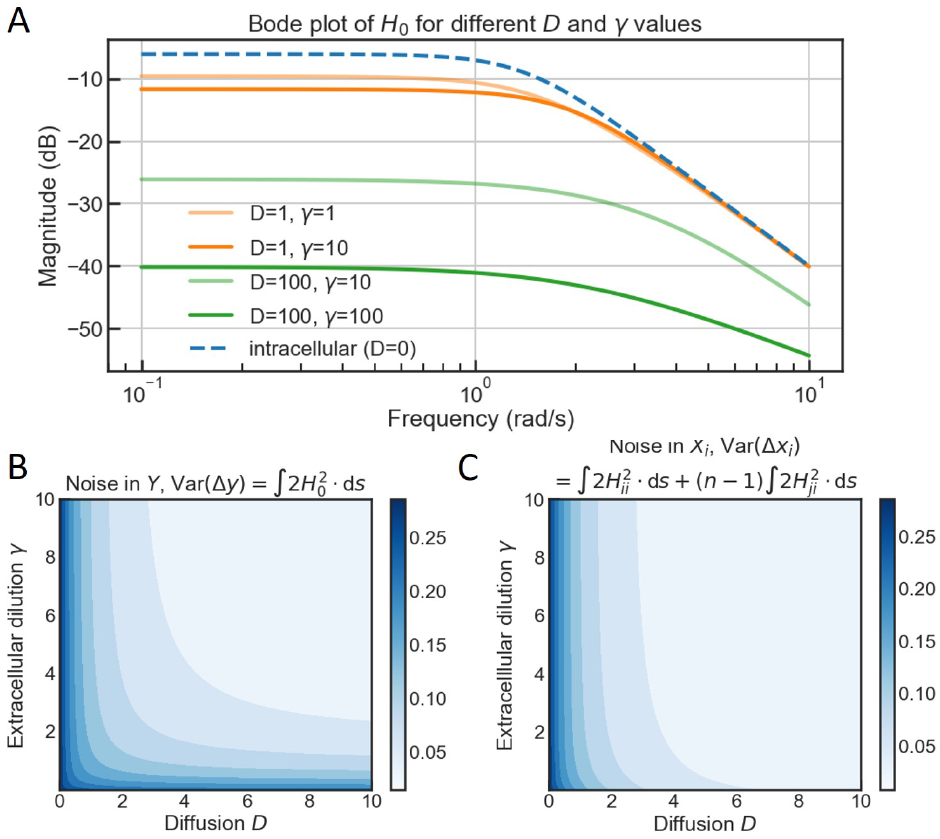
Numerical analysis of transfer functions in the frequency domain and expression variance with dynamic measurement noise Δ*β*_*i*_(*t*). (A) Bode plots (magnitude) of transfer functions *H*_0_ of open loop (dashed black), intracellular feedback (blue dashed) and signal-based population feedback (orange and green) with different parameters. (B) Dependence of variance of noise in *Y* on diffusion *D* and extracellular dilution *γ*. (C) Dependence of variance of noise in *X*_*i*_ on diffusion *D* and extracellular dilution *γ*.

This conclusion is particularly intuitive, since measurement noise arises from the fluctuations in the signaling molecule production (the controller), rather than directly from the targe gene expression (the process). Therefore, the averaging effects of cell-cell signaling help smooth out variations in the controller, leading to a more robust control output to the process.

## V. Independent control of mean and variance of target gene expression

Building upon the analysis of noise regulation at both single-cell and population levels for process and measurement noise sources, we observe that the mean and variance of target gene expression can be controlled by tuning different parameters.

In previous sections, we set 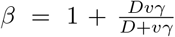 to ensure identical steady-state solutions for 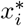 and *y*^*^ across different systems for comparison. In practice, the parameters *β* and *γ* can be independently tuned, by altering the production rate and extracellular dilution rate of signaling molecules, respectively. For simplicity, we define a lumped parameter 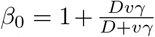. We can tune *β*_0_ by varying the extracellular dilution rate *γ*. Based on Eq (4), the production rate *β* can be expressed as 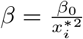.

For static process noise Δ*α*_*i*_, we can derive 𝔼 [*x*_*i*_] and Var[*x*_*i*_] based on Eq (17) and (18). Under the fast diffusion condition where *D* ≫ 1, *vγ*, we obtain:

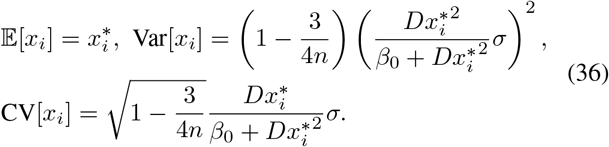

By tuning the extracellular dilution rate *γ*, we can adjust *β*_0_ and control the variance and CV[*x*_*i*_] independently of its mean.

At the population level, we have:

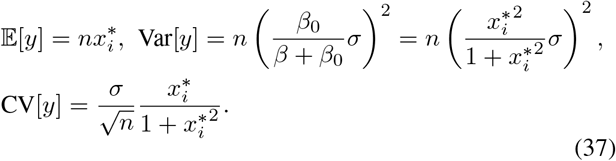

The result shows that the variance and CV[*y*] cannot be tuned by any other parameters except the population density *n*, given a desired mean value. Therefore, in a population of *n* cells, the population-level mean and variance are coupled with no feasible designs for independent control.

Similarly, for static measurement noise Δ*β*, we can derive the mean and variance at single-cell level based on Eq (33) and (34). Consider the fast diffusion condition where *D* ≫ 1, *vγ*, we obtain:

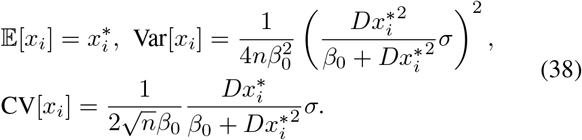

At population level, we have:

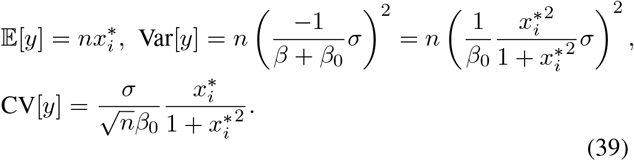

In this case, mean and variance at single-cell and population levels can be independently controlled by tuning *β* and *γ*.

## VI. Conclusion

In multicellular systems, cell-cell signaling plays a vital role in achieving robust population-level behaviors. It synchronizes individual cellular dynamics and enables coherent population responses through averaging effects [10], [17], [18]. Furthermore, when combined with negative feedback, cell-cell signaling provides an additional layer of control for stabilizing and fine-tuning population-level functionalities [13], [19], [20].

Our analysis reveals that the interplay between cell-cell signaling and negative feedback may exhibit contrasting effects on noise regulation and disturbance rejection at single-cell and population levels. For measurement noise arising from fluctuations in cell-cell signaling dynamics, both single-cell and population-level noise are effectively attenuated under strong averaging effects. These effects are enhanced by fast signal diffusion, efficient extracellular dilution, large population size, and a sufficiently high extracellular volume. In contrast, for process noise, cell-cell signaling primarily reduces noise at the population level, especially dynamic noise under conditions of fast diffusion and extracellular dilution. However, this mechanism fails to attenuate noise at the single-cell level, leading to increased cell-to-cell variability compared to systems with intracellular-only negative feedback.

The inconsistent noise regulation properties between singlecell and population levels suggest that cell-to-cell variability and population-level robustness are inherently coupled, making the independent control of these two layers challenging and potentially suboptimal.In fact, phenotype variability has been shown to enhance population robustness, enabling multicellular systems to adapt and survive in fluctuating environments [21], [22]. Thus, population-level stability and robustness can emerge as a result of single-cell variability. In heterogeneous populations, cell-cell signaling can propagate this variability, enabling collective adaptation to extrinsic disturbances. Moreover, the fundamental limitations of negative feedback, such as the requirement for fast and abundant production of regulatory molecules, impose significant burdens on individual cells [23]. In such cases, cell-cell signaling alleviates the burden of high-gain feedback by redistributing the regulatory load across the entire population. Our study primarily focuses on scenarios where noise is independent and identically distributed (i.i.d.) across all cells. In other scenarios, a cell population may encounter non-uniform extrinsic disturbances. By enabling distributed compensatory production, cell-cell signaling ensures that minor adjustments across many cells collectively buffer against noise, thereby preventing the need for disproportionately high production in a few affected cells.

From a design perspective, our analysis suggests that specific parameters, such as signaling molecule production rate and extracellular dilution rate, can be fine-tuned to independently control the mean and variance of target gene expression under different noise conditions. However, the coupling between mean and variance for process noise at the population level highlights a fundamental limitation of population-level negative feedback in noise attenuation. Additionally, our frequency-domain analysis elucidates how different noise sources propagate through population feedback systems, providing insights for optimizing noise reduction in multicellular systems. These findings pave the way for the systematic design of population-level controllers by employing loop-shaping techniques, such as transfer function analysis [24]. Furthermore, integrating feedforward control with feed-back mechanisms can enhance noise attenuation and improve system robustness. These strategies offer a comprehensive framework for designing synthetic circuits tailored to achieve robust and tunable gene expression in multicellular systems.

## Appendix

### A. Mean and variance of signaling molecule concentrations

Based on the linearized model of the signal-based population feedback system with process noise Δ*α*_*i*_ in Eq (11), we can derive the following expression for single-cell level noise in signaling molecule concentrations Δ*s*_*i*_:

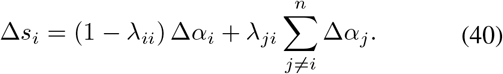

The characteristics of noise in *S*_*i*_ are:

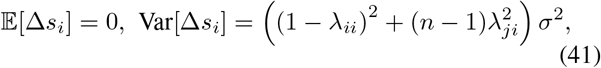

depending on parameters *D, v, γ* and *n*. Now we consider a strong averaging effect from cell-cell signaling (*D* and *n* are large), then we can approximate:

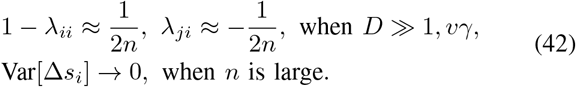

#### B. Linearized models for measurement noise

For signal-based population feedback system, the linearized model given measurement noise Δ*β*_*i*_ is as follows:

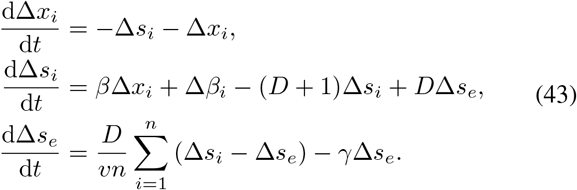

#### C. Derivation of transfer functions for measurement noise

We perform a Fourier transformation of the population feed-back system as described in Eq (43). The resulting population-level noise is:

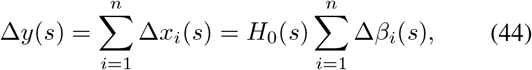

where the transfer function *H*_0_(*s*) is given by:

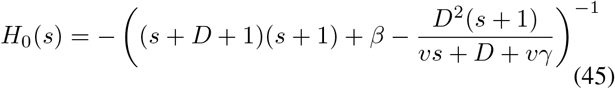

This describes how all measurement noise 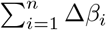 propagates to the population expression. At single-cell level, the expression noise is given by:

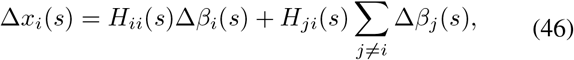

with the corresponding transfer functions *H*_*ii*_(*s*) and *H*_*ji*_(*s*):

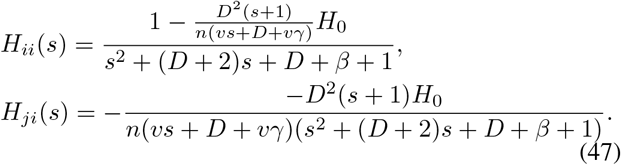

*H*_*ii*_(*s*) and *H*_*ji*_(*s*) describe how measurement noise from individual cells and other neighboring cells affect *X* expression in a single cell.

## Acknowledgment

The authors would like to thank Fangzhou Xiao, Rongrong Du, Christian Cuba Samaniego and Frank Britto Bisso for their insightful discussion and feedback.

## Notes

### Competing Interest Statement

The authors have declared no competing interest.

